# Introduction of two prolines and removal of the polybasic cleavage site leads to optimal efficacy of a recombinant spike based SARS-CoV-2 vaccine in the mouse model

**DOI:** 10.1101/2020.09.16.300970

**Authors:** Fatima Amanat, Shirin Strohmeier, Raveen Rathnasinghe, Michael Schotsaert, Lynda Coughlan, Adolfo García-Sastre, Florian Krammer

## Abstract

The spike protein of severe acute respiratory syndrome coronavirus 2 (SARS-CoV-2) has been identified as the prime target for vaccine development. The spike protein mediates both binding to host cells and membrane fusion and is also so far the only known viral target of neutralizing antibodies. Coronavirus spike proteins are large trimers that are relatively instable, a feature that might be enhanced by the presence of a polybasic cleavage site in the SARS-CoV-2 spike. Exchange of K986 and V987 to prolines has been shown to stabilize the trimers of SARS-CoV-1 and the Middle Eastern respiratory syndrome coronavirus spikes. Here, we test multiple versions of a soluble spike protein for their immunogenicity and protective effect against SARS-CoV-2 challenge in a mouse model that transiently expresses human angiotensin converting enzyme 2 via adenovirus transduction. Variants tested include spike protein with a deleted polybasic cleavage site, the proline mutations, a combination thereof, as well as the wild type protein. While all versions of the protein were able to induce neutralizing antibodies, only the antigen with both a deleted cleavage site and the PP mutations completely protected from challenge in this mouse model.

**Importance:** A vaccine for SARS-CoV-2 is urgently needed. A better understanding of antigen design and attributes that vaccine candidates need to have to induce protective immunity is of high importance. The data presented here validates the choice of antigens that contain the PP mutation and suggests that deletion of the polybasic cleavage site could lead to a further optimized design.

## Introduction

Severe acute respiratory syndrome coronavirus 2 (SARS-CoV-2) emerged in late 2019 in China and has since then caused a coronavirus disease 2019 (COVID-19) pandemic (1–3). Vaccines are an urgently needed countermeasure to the virus. Vaccine candidates have been moved at unprecedented speed through the pipeline with first Phase III trials already taking place in summer 2020, only half a year after discovery of the virus sequence. From studies on SARS-CoV-1 and the Middle Eastern respiratory syndrome CoV (MERS-CoV), it was clear that the spike protein of the virus is the best target for vaccine development (4–6). Most coronaviruses (CoVs) only have one large surface glycoprotein (a minority also have a hemagglutinin-esterase) that is used by the virus to attach to the host cell and trigger fusion of viral and cellular membranes. The spike protein of SARS-CoV-2, like the one of SARS-CoV-1, binds to human angiotensin receptor 2 (ACE2) (7–9). In order to be able to trigger fusion, the spike protein has to be cleaved into the S1 and S2 subunit (10–12). Additionally, a site in S2 (S2’) that has to be cleaved to activate the fusion machinery has been reported as well (13). While the spike of SARS-CoV-1 contains a single basic amino acid at the cleavage site between S1 and S2, SARS-CoV-2 has a polybasic motif that can be activated by furin-like proteases (10–12), analogous to the hemagglutinin (HA) of highly pathogenic H5 and H7 avian influenza viruses. In addition, it has been reported that the activated spike protein of CoVs is relatively instable and multiple conformations might exist of which not all may present neutralizing epitopes to the immune system. For SARS-CoV-1 and MERS-CoV stabilizing mutations – a pair of prolines replacing K986 and V987 in S2 – have been described (14) and a beneficial effect on stability has also been shown for SARS-CoV-2 (9). Here, we set out to investigate if including these stabilizing mutations, removing the cleavage site between S1 and S2 or combining the two strategies to stabilize the spike would increase its immunogenicity and protective effect in a mouse model that transiently expressed hACE2 via adenovirus transduction (15). This information is important since it can help to optimize vaccine candidates, especially improved versions of vaccines that might be licensed at a later point in time.

## Results

### Construct design and recombinant protein expression

The sequence based on the S gene of SARS-CoV-2 strain Wuhan-1 was initially codon optimized for mammalian cell expression. The wild type signal peptide and ectodomain (amino acid 1-1213) were fused to a T4 foldon trimerization domain followed by a hexa-histidine tag to facilitate purification. This construct was termed wild type (WT). Additional constructs were generated including one in which the polybasic cleavage site (RRAR) was replaced by a single alanine (termed ΔCS), one in which K986 and V987 in the S2 subunit were mutated to prolines (PP) and one in which both modifications were combined (ΔCS-PP) (**Figure 1A-C**). The proteins were then expressed in a baculovirus expression system and purified. At first inspection by sodium dodecyl sulfate polyacrylamide gel electrophoresis (SDS-PAGE) and Coomassie staining, all four constructs appeared similar with a major clean band at approximately 180kDa (**Figure 1E**). When a Western blot was performed, additional bands were detected in the lanes with the WT, PP and ΔCS-PP constructs, suggesting cleavage of a fraction of the protein. For WT, the most prominent detected smaller band ran at 80 kDa, was visualized with an antibody recognizing the C-terminal hexa-histidine tag and likely represents S2 (**Figure 1F**). The two constructs containing the PP mutations also produced an additional band at approximately 40 kDa (**Figure 1E**), potentially representing a fragment downstream of S2’. While in general these bands were invisible on an SDS PAGE and therefore are likely only representing a tiny fraction of the purified spike protein, they might indicate vulnerability to proteolytic digest of the antigen *in vivo*. All constructs were also recognized in a similar manner by mAb CR3022 (16, 17), an antibody that binds to the RBD (**Figure 1F**).

**Figure 1.**
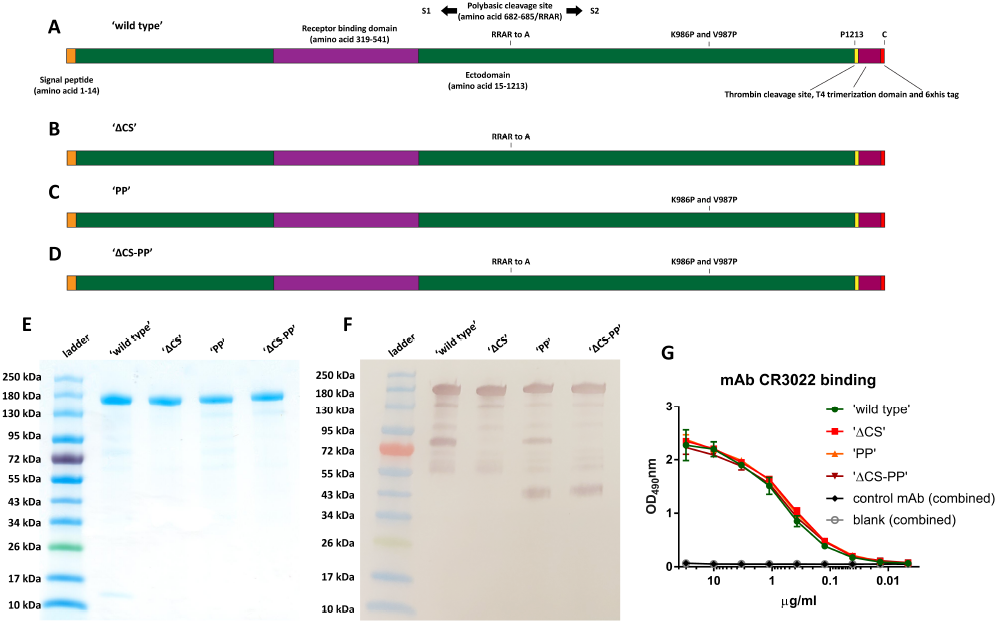
Spike construct design and protein characterization. (A-D) described the wild type, ΔCS, PP and ΔCS-PP constructs used in this study. (B) shows the four antigens on a SDS-PAGE stained with Coomassie blue, while (C) shows the same protein on a Western blot developed with an antibody to the C-terminal hexahistidine tag. While all four proteins are detected as clean, single bands on the SDS-PAGE, the Western blot reveals a small fraction of degradation products at approximately 80 kDa for the wild type and PP variants and of approximately 40 kDa for the PP and ΔCS-PP constructs. (D) shows binding of mAb CR3022 to the constructs in ELISA. Data for the negative control mAb and the blank were combined for the different substrates.

### All versions of the recombinant spike protein induce robust immune responses in mice

To test the immunogenicity of the four spike constructs, all proteins were used in a simple prime-boost study in mice (**Figure 2A**). Animals were injected intramuscularly (i.m.) with 3μg of spike protein adjuvanted with AddaVax (a generic version of the oil-in-water adjuvant MF59) twice in a 3 week interval. A control group received an irrelevant immunogen, recombinant influenza virus hemagglutinin (HA), also expressed in insect cells, with AddaVax. Mice were bled three weeks after the prime and four weeks after the boost to assess the immune response that they mounted to the vaccine (**Figure 2B**). To determine antibody levels to the RBD, we performed enzyme-linked immunosorbent assays against recombinant, mammalian cell expressed RBD (18, 19). All animals made anti-RBD responses after the prime but they were higher in the ΔCS and ΔCS-PP groups than in the WT or PP groups (**Figure 2C**). The booster dose increased antibodies to the RBD significantly but the same pattern persisted (**Figure 2D**). Interestingly, the ΔCS-PP group showed very homogenous responses compared to the other groups were there was more spread between the animals. In addition, we also performed cell-based ELISAs with Vero cells infected with SARS-CoV-2 as target. While all groups showed good reactivity, a similar pattern emerged in which ΔCS and ΔCS-PP groups showed higher reactivity than WT and PP groups (**Figure 2E**). Finally, we performed microneutralization assays with authentic SARS-CoV-2 (20). Here, the WT, PP and ΔCS groups showed similar levels of neutralization while the ΔCS-PP group animals had higher serum neutralization titers (**Figure. 2F**).

**Figure 2.**
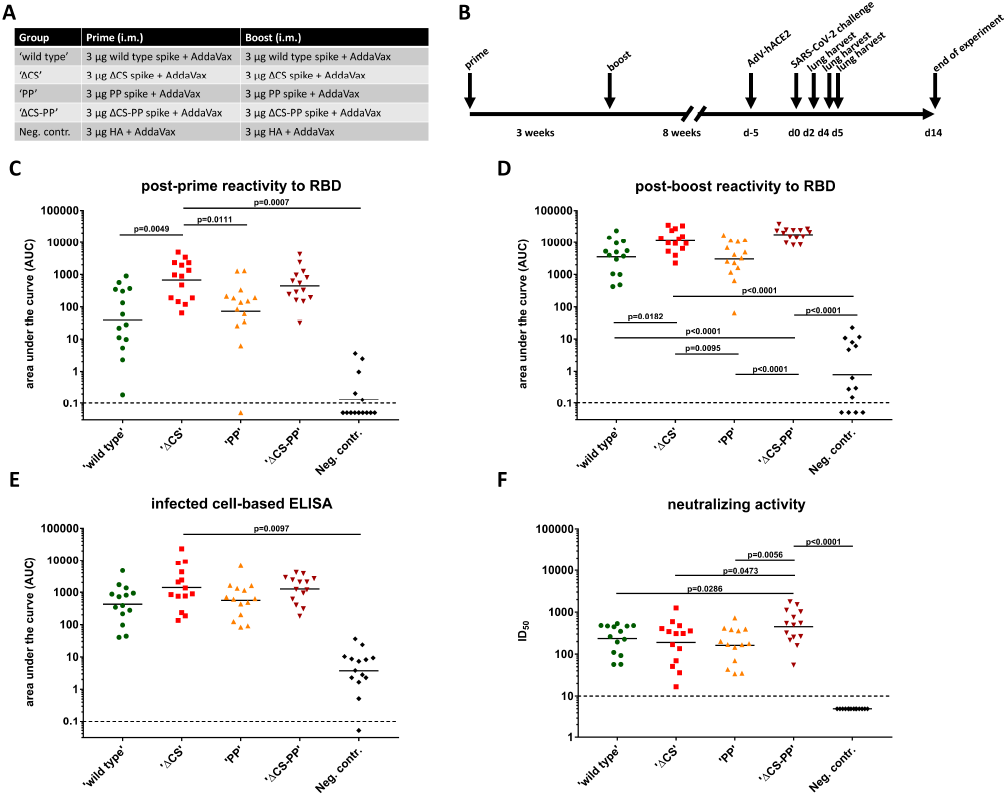
Immunogenicity of different spike variants in the mouse model. (A) shows the vaccination regimen used for the five groups of mice and (B) shows the timeline. Animals were bled 3-weeks post prime (C) and 4 weeks post-boost (D) and antibody levels to a mammalian-cell expressed RBD were measured. Post-boost sera were also tested in cell-based ELISAs on cells infected with authentic SARS-CoV-2. Finally, post-boost sera were tested in a microneutralization assay against SARS-CoV-2.

### Vaccination with recombinant S protein variants protects mice from challenge with SARS-CoV-2

In order to perform challenge studies, mice were sensitized to infection with SARS-CoV-2 by intranasal (i.n.) transduction with an adenovirus expressing hACE2 (AdV-hACE2), using a treatment regimen described previously (**Figure 2A**) (15, 21, 22). They were then challenged with 10^5^ plaque forming units (PFU) of SARS-CoV-2 and monitored for weight loss and mortality for 14 days. Additional animals were euthanized on day 2 and day 4 to harvest lungs for histopathological assessment and immunohistochemistry, and on day 2 and day 5 to measure virus titers in the lung. After challenge, all groups lost weight trending with the negative control group (irrelevant HA protein vaccination), except for the ΔCS-PP group which displayed minimal weight loss (**Figure 3A**). Only on days 4-6 the WT, PP and ΔCS groups showed a trend towards less weight loss then the control group. However, all animals recovered and by day 14 and no mortality was observed. Lung titers on day 2 suggested low virus replication in the WT, PP and ΔCS groups with some animals having no detectable virus and no presence of replication competent virus in the ΔCS-PP (**Figure 3B**). Two of the control animals showed high virus replication while virus could not be recovered from the third animal. No virus could be detected in any of the vaccinated groups on day 5 while all three controls still had detectable virus in the 10^4^ to 10^5^ range (**Figure 3C**).

**Figure 3.**
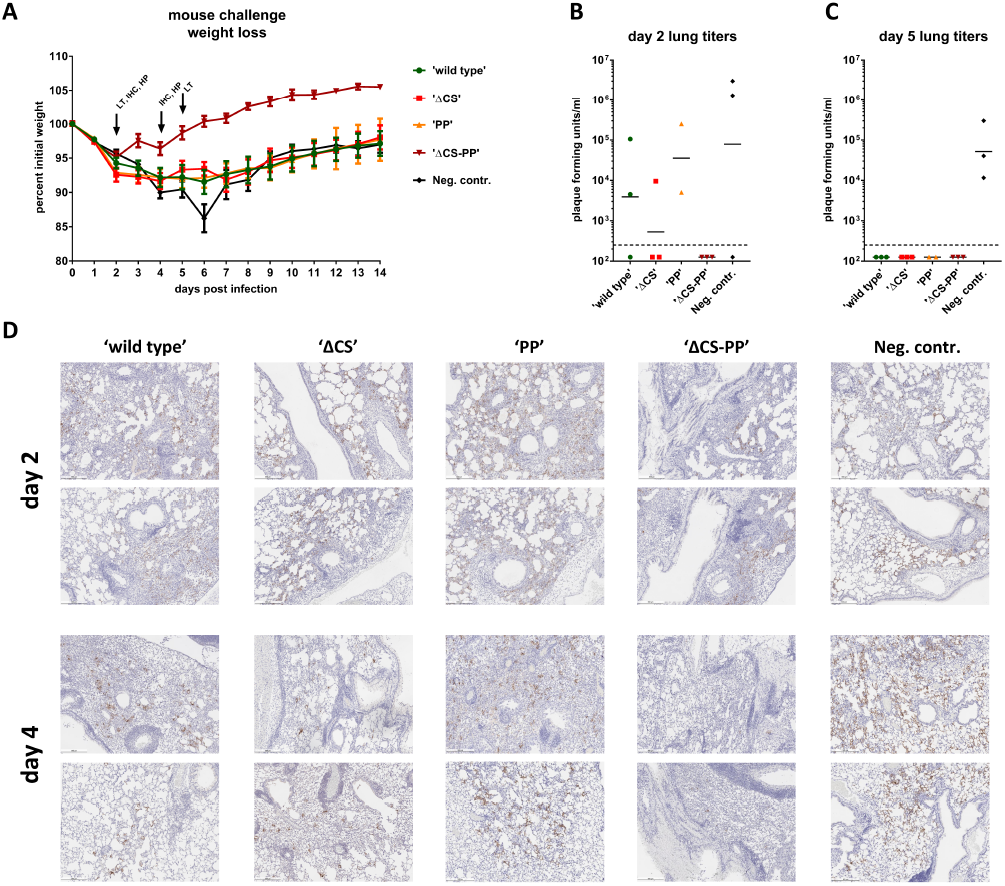
Challenge of mice with SARS-CoV-2. Animals sensitized by transient expression of hACE2 via adenovirus transduction were challenged with 10^5^ PFU or SARS-CoV-2 and weight loss was monitored over a period of 14 days (A). (B) and (C) shows day 2 and day 5 lung titers respectively, while (D) shows lung immunohistochemistry staining for SARS-CoV-2 nucleoprotein on days 2 and 4 post challenge. Representative images from two animals each are shown at 5-fold magnification. Scale bar = 500 um.

### Lung immunohistochemistry and pathology

Lungs were harvested on days 2 and 4 post challenge. Samples from both days were used for immunohistochemistry to detect viral nucleoprotein antigen. Viral antigen was detectable in all groups on day 2 as well as day 4 post infection (**Figure 3D**). However, the ΔCS-PP group showed very few positive cells, especially on day 4 while antigen was detected more widespread in all other groups. These results correlate well with the viral lung titers shown above. The samples were also hematoxylin and eosin (H&E) stained and scored for lung pathology by a qualified veterinary pathologist using a composite score with a maximum value of 24 (**Figure 4A and C**). At D2 post-infection with SARS-CoV-2, all mice were determined to exhibit histopathological lesions typical of interstitial pneumonia, with more severe alveolar inflammation in the WT group. Alveolar congestion and edema were also more pronounced in S vaccinated groups as compared with the irrelevant control HA immunogen. At this time-point, the overall pathology score was lowest for the irrelevant HA control group, followed by ΔCS-PP<PP<ΔCS<wild type (**Figure 4A**). On day 4 all groups showed mild to moderate pathology scores, reduced in severity as compared with D2. Observations included perivascular, bronchial and alveolar inflammation, as well as mild to moderate congestion or edema. Scores were slightly higher in vaccinated than control animals which may reflect the infiltration of CoV-2 antigen-specific immune cells into the lung, which would be absent in the irrelevant HA immunized control mice (**Figure 4C and D**).

**Figure 4:**
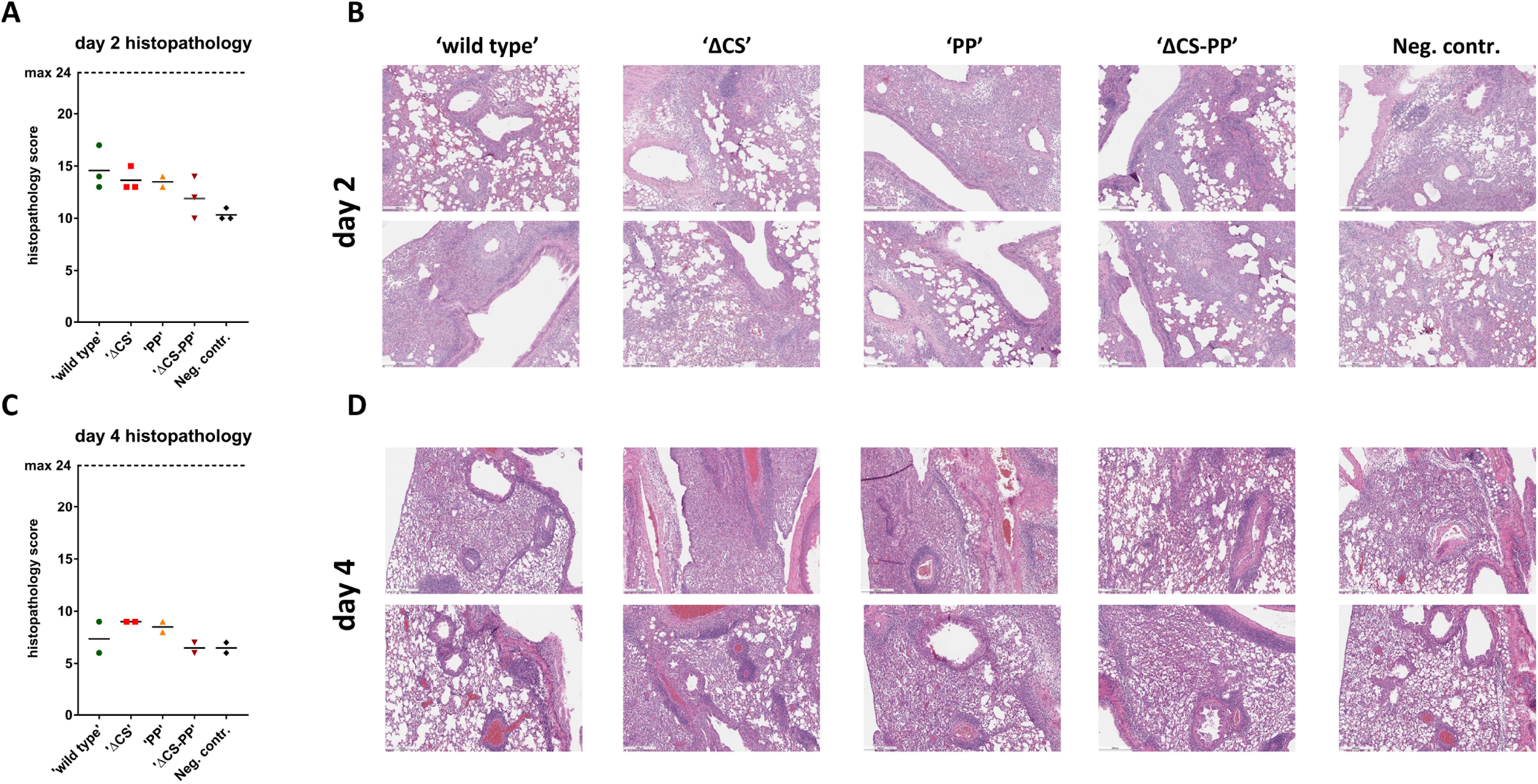
Lung pathology. (A) shows a histopathological composite score for animals on day 2 post infection, (B) shows representative H&E stained tissue images from 2 animals per group. (C) and (D) show the same but for day 4 post challenge. Scale bar = 500 um.

## Discussion

The spike protein of SARS-CoV-2 has been selected early on as a target for vaccine development, based on experience with SARS-CoV-1 and MERS CoV (6). The coronavirus spike protein is known to be relatively labile, and in addition to this inherent property the SARS-CoV-2 spike also contains a polybasic cleavage site between S1 and S2. Work on SARS-CoV-1 and MERS CoV had shown that introducing two prolines in positions 986 and 987 (SARS-CoV-2 numbering) improves stability and expression (14). In addition, removal of polybasic cleavage sites has been shown to stabilize hemagglutinin (HA) proteins of highly pathogenic influenza viruses. In this study, we tested different versions of the protein either lacking the polybasic cleavage site or including the stabilizing PP mutations or both. While vaccination with all constructs induced neutralizing antibodies and led to control of virus replication in the lung, we observed notable differences. Removing the polybasic cleavage side did increase the humoral immune response in ELISAs. Since we did not observe cleavage of the majority of protein when purified (although some cleavage could be observed), even with the polybasic cleavage site present, we speculate that removal of the site might make the protein more stable *in vivo* post vaccination. Longer stability could lead to stronger and potentially more uniform immune responses. The combination of deleting the polybasic cleavage site plus introducing the PP mutations performed best, also in terms of protection of mice from weight loss. It is important to note that all versions of the protein tested had a third stabilizing element present, which is a trimerization domain. This trimerization domain might have also increased stability and immunogenicity.

Current leading vaccine candidates in clinical trials include virus vectored and mRNA vaccines. The ChAdOx based vaccine candidate that is developed by AstraZeneca is using a wild type version of the spike protein (23), while Moderna’s mRNA vaccine is based on a spike construct that includes the PP mutations but features a wild type cleavage site (24). It is currently unclear, if addition of the modifications shown here to enhance immunogenicity of recombinant protein spike antigens would also enhance immunogenicity of these constructs. However, it might be worth testing if these vaccine candidates can be improved by our strategy as well. Of note, ones study in non-human primates with adenovirus 26-vectored vaccine candidates expressing different versions of the spike protein also showed that a ΔCS-PP (although including the transmembrane domain) performed best and this candidate is now moving forward into clinical trials (25). Similarly, Novavax is using a recombinant spike construct that features ΔCS-PP and, when adjuvanted, induced high neutralization titers in humans in a Phase I clinical trial (26).

While vaccination with all constructs led to various degrees of control of virus replication, histopathology scores, especially on day 2 after challenge were above those of the negative controls animals. We do not believe that this is a signal of enhanced disease as it has been observed in some studies for SARS-CoV-2 but the hallmark of an antigen-specific immune response. This is also evidenced by significantly reduced weight loss in the ΔCS-PP group as well as complete control of virus replication despite having increased lung histopathology scores. However, future studies with recombinant protein vaccines that are routed for clinical testing, as outlined below, will need to assess this increase in lung pathology in more detail.

Recombinant protein vaccines including the spike ectodomain (27, 28), membrane extracted spike (29) as well as S1 (30) and RBD (31) have been tested for SARS-CoV-1 and several studies show good efficacy against challenge in animal models. It is, therefore, not surprising that similar constructs for SARS-CoV-2 also provided protection. While our goal was not vaccine development but studying the effect of stabilizing elements on the immunogenicity of the spike protein, Sanofi Pasteur has announced the development of a recombinant protein based SARS-CoV-2 vaccine and a second recombinant protein candidate is currently being developed by Seqirus. Our data shows that this approach could be effective.

## Materials and methods

### Cells and viruses

Vero.E6 cells (ATCC CRL-1586-clone E6) were maintained in culture using Dulbecco’s Modified Eagle Medium (DMEM; Gibco) which was supplemented with Antibiotic-Antimycotic (100 U/ml penicillin-100 μg/ml streptomycin-0.25 ug/ml Amphotericin B) (Gibco; 15240062) and 10% fetal bovine serum (FBS; Corning). SARS-CoV-2 (isolate USA-WA1/2020 BEI Resources, NR-52281) was grown in Vero.E6 cells as previously described and was used for the *in vivo* challenge (20). A viral seed stock for a non-replicating human adenovirus type-5 (HAdV-C5) vector expressing the human ACE2 receptor was obtained from the Iowa Viral Vector Core Facility. High titer Ad-hACE2 stocks were amplified in TRex™-293 cells, purified by CsCl ultracentrifugation and infectious titers determined by tissue-culture infectious dose-50 (TCID_50_), adjusting for plaque forming unit (PFU) titers using the Kärber statistical method, as described previously (32).

### Recombinant proteins

All recombinant proteins were expressed and purified using the baculovirus expression system, as previously described (18, 33, 34). Different versions of the spike protein of SARS-CoV-2 (GenBank: MN908947.3) were expressed to assess immunogenicity. PP indicates that two stabilizing prolines were induced at K986 and K987. ΔCS indicates that the cleavage site of the spike protein was removed by deletion of the arginine residues (RRAR to just A). The HA was also produced in the baculovirus expression system similar to the spike variants.

### SDS-PAGE and Western blot

One ug of each respective protein was mixed at a 1:1 ratio with 2X Laemmli buffer (Bio-Rad) which was supplemented with 2% β-mercaptoethanol (Fisher Scientific). The samples were heated at 90°C for 10 minutes and loaded onto a 4-20% precast polyacrylamide gel (BioRad). The gel was stained with SimplyBlue SafeStain (Invitrogen) for 1 hour and then de-stained with water for a few hours. For Western blot, the same process was used as mentioned above. After the gel was run, the gel was transferred onto a nitrocellulose membrane, as described previously (33). The membrane was blocked with phosphate buffered saline (PBS; Gibco) containing 3% non-fat milk (AmericanBio, catalog# AB10109-01000) for an hour at room temperature on an orbital shaker. Next, primary antibody was prepared in PBS containing 1% non-fat milk using anti-hexahistidine antibody (Takara Bio, catalog #631212) at a dilution of 1:3000. The membrane was stained with primary antibody solution for 1 hour at room temperature. The membrane was washed thrice with PBS containing 0.1% Tween-20 (PBS-T; Fisher Scientific). The secondary solution was prepared with 1% non-fat milk in PBS-T using anti-mouse IgG (whole molecule)–alkaline phosphatase (AP) antibody produced in goat (Sigma-Aldrich) at a dilution of 1:3,000. The membrane was developed using an AP conjugate substrate kit, catalog no. 1706432 (Bio-Rad).

### ELISA

Ninety-six well plates (Immulon 4 HBX; Thermo Fisher Scientific) were coated with recombinant RBD at a concentration of 2 ug/ml with 50μl/well overnight. The RBD protein was produced in 293F cells and purified using Ni-NTA resin and this procedure has been described in detail earlier (19). The next morning, coating solution was removed and plates were blocked with 100μls of 3% non-fat milk (AmericanBio, catalog# AB10109-01000) prepared in PBS-T for 1 hour at room temperature (RT). Serum samples from vaccinated mice were tested on the ELISA starting at a dilution of 1:50 and three-fold subsequent dilutions were performed. Serum samples were prepared in PBS-T containing 1% non-fat milk and the plates were incubated with the serum samples for 2 hours at RT. Next, plates were washed with 200μls of PBS-T thrice. Anti-mouse IgG conjugated to horseradish peroxidase (Rockland, catalog# 610-4302) was used at a concentration of 1:3000 in PBS-T with 1% non-fat milk and 100 μl was added to each well for 1 hour at RT. Plates were then washed again with 200 uls of PBS-T and patted dry on paper towel. Developing solution was prepared in sterile water (WFI, Gibco) using SIGMAFAST OPD (o-phenylenediamine dihydrochloride; Sigma-Aldrich, catalog# P9187) and 100 μls was added to each well for a total of 10 mins. Next, the reaction was stopped with 50 μls of 3M hydrochloric acid and absorbance was measured at 490 nm (OD_490_) using a Synergy 4 (BioTek) plate reader. Data was analyzed using GraphPad Prism 7 and are under the curve (AUC) values were measured and graphed (18). An AUC 0f 0.05 was assigned to negative values for data analysis purposes.

To perform an ELISA on infected cells, Vero.E6 cells were seeded at 20,000 cells per well in a 96-well cell culture plate a day before and infected at a multiplicity of infection of 0.1 for 24 hours with SARS-CoV-2 (isolate USA-WA1/2020 BEI Resources, NR-52281). The cells were fixed with 10% formaldehyde (Polysciences) for 24 hours after which the ELISA procedure mentioned above was performed using serum from each vaccinated animal.

### Mouse vaccinations and challenge

All animal procedures were performed by adhering to the Institutional Animal Care and Use Committee (IACUC) guidelines. Six to eight week old, female, BALB/c mice (Jackson Laboratories) were immunized intramuscularly with 3 μg of recombinant protein per mouse with an adjuvant, AddaVax (Invivogen) in a volume of 50 ul. Three weeks later, mice were again immunized, via intramuscular route, with 3 μg of each respective protein with adjuvant. Mice were bled 3 weeks after the prime regimen and were also bled 4 weeks after the boost regimen. Another four weeks later, 2.5×10^8^ PFU/mouse of AdV-hACE2 was administered intranasally to each mouse in a final volume of 50μL sterile PBS. Adhering to institutional guidelines, a mixture containing 0.15 mg/kg ketamine and 0.03 mg/kg xylazine in water was used as anesthesia for mouse experiments and intranasal infection was performed under anesthesia.

Five days post administration of the AdV-hACE2, mice were infected with 10^5^ PFUs of SARS-CoV-2. On day 2 and day 5, mice were sacrificed using humane methods and the whole lung was dissected from each mouse. Mice were sacrificed for measuring viral titers in the lung as well as to see pathological changes in the lungs. Lungs were homogenized using BeadBlaster 24 (Benchmark) homogenizer after which the supernatant was clarified by centrifugation at 14,000 g for 10 mins. Experimental design was adapted from earlier reported work (15, 35). The remaining mice were weight daily for 14 days.

### Micro-neutralization assays

We used a very detailed protocol that we published earlier for measuring neutralizing antibody in serum samples (18, 20). Briefly, Vero.E6 cells were seeded at a density of 20,000 cells per well in a 96-well cell culture plate. Serum samples were heat-inactivated for 1 hour at 56 C. Serial dilutions starting at 1:10 were prepared in 1X (minimal essential medium; MEM) supplemented with 1% FBS. The remaining steps of the assay were performed in a BSL3 facility. Six-hundred TCID_50_ of virus in 80μls was added to 80 uls of each serum dilution. Serum-virus mixture was incubated at room temperature for 1 hour. After 1 hour, media from the cells was removed and 120 μls of serum-virus mixture was added onto the cells. The cells were incubated for 1 hour in a 37°C incubator. After 1 hour, all of the serum-virus mixture was removed. One hundred uls of each corresponding serum dilution was added onto the cells and 100 uls of 1X MEM was added to the cells as well. The cells were incubated at 37°C for 2 days. After 2 days, cells were fixed with 10% formaldehyde (Polysciences). The next day, cells were stained with an anti-nuceloprotein antibody (ThermoFisher; Catalog # PA5-81794) according to our published protocol (20). The 50% inhibitory dilution (ID_50_) for each serum was calculated and the data was graphed. Negative samples were reported as half of the limit of detection (ID_50_ of 5).

### Plaque assays

Four-hundred thousand Vero.E6 cells were plated the day before the plaque assay was performed. All assays using SARS-CoV-2 were performed in the BSL3 following institutional guidelines. To assess viral titer in the lung, plaque assays were performed using lung homogenates. Dilutions of lung homogenates were prepared starting from 10^−1^ to 10^−6^ in 1X MEM supplemented with 2% FBS. Media was removed from cells and each dilution was added to the cells. The cells were incubated in a humidified incubator at 37°C for 1 hour. Next, the virus was removed, and cells were overlaid 2X MEM supplemented with 2% oxoid agar (final concentration of 0.7%) as well as 4% FBS. The cells were incubated at 37°C for 72 hours after which cells were fixed with 1 ml of 10% formaldehyde (Polysciences) was added for 24 hours to ensure inactivation of virus. Crystal violet was used to visualize the plaques. Only 5 or more plaques were counted and the limit of detection was 250.

### Histology and immunohistochemistry

Mice were subjected to terminal anesthesia and euthanasia performed by exsanguination of the femoral artery before lungs were flushed/inflated with 10% formaldehyde by injecting a 19 gauge needle through the trachea on day 4 for immunohistochemistry. Fixed lungs were sent to a commercial company, Histowiz for paraffin embedding, tissue analysis and scoring by an independent veterinary pathologist. Hematoxylin and eosin (H&E) and immunohistochemistry (IHC) staining was performed. IHC staining was performed using an anti-SARS-CoV nucleoprotein antibody (Novus Biologicals cat. NB100-56576). Histology and IHC for day 2 samples was performed on only half of the lung which was dissected and cut in half from sacrificed mice. The other half of the lung was used for quantification of virus, as mentioned above.

Scores were assigned by the pathologist based on six parameters: perivascular inflammation, bronchial/bronchiolar epithelial degeneration/necrosis, bronchial/bronchiolar inflammation, intraluminal debris, alveolar inflammation and congestion/edema. A 5-point scoring system was used ranging from 0-4, with 0 indicating no epithelial degeneration/necrosis and inflammation while 4 indicating severe epithelial degeneration/necrosis and inflammation.

### Statistics

Statistical analysis was performed in Graphpad Prism using one-way ANOVA corrected for multiple comparisons.

## Acknowledgments

We thank Randy A. Albrecht for oversight of the conventional BSL3 biocontainment facility. This work was partially supported by the NIAID Centers of Excellence for Influenza Research and Surveillance (CEIRS) contract HHSN272201400008C (FK, AG-S), Collaborative Influenza Vaccine Innovation Centers (CIVIC) contract 75N93019C00051 (FK, AG-S),, NIAID R21AI157606 (L.C), a supplement to NIAD adjuvant contract HHSN272201800048C (FK, AG-S) and the generous support of the JPB foundation, the Open Philanthropy Project (#2020-215611 to AG-S) and other philanthropic donations. We thank Susan Stamnes, Kaylee Murphy (Iowa Viral Vector Core) and Dr. Paul B. McCray Jnr (University of Iowa) for making the AdV-hACE2 virus rapidly available to us.

## Data availability

Raw data is available from the corresponding author upon reasonable request.

